## supplemental files for "Progressive remote memory decline coincides with parvalbumin interneuron hyperexcitability and enhanced inhibition of cortical engram cells in a mouse model of Alzheimer’s disease"

Supplemental figures

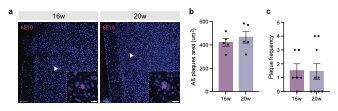

**Sup. Fig. 1:** **Amyloid beta plaque load in mPFC at 16 and 20 weeks of age does not differ.** **a.** Representative images at 12w (left) and 16w (right) in APP/PS1 mice showing DAPI staining (blue) and 6E10 amyloid plaque (magenta). Scale bars, main image: 100 µm; insets: 25 µm. **b.** No difference in average amyloid plaque area (µm^2^) in the mPFC between APP/PS1 mice at 16w and 20w. Unpaired *t*-test: *t*_20_ = 0.84, *p* = 0.41. **c**. No difference in the plaque frequency in the mPFC between APP/PS1 mice at 16w and 20w. Mann Whitney test: *U* = 24, *p* = 0.39. 16w (*n* = 6); 20w (*n* = 9). Graphs show mean ± s.e.m.

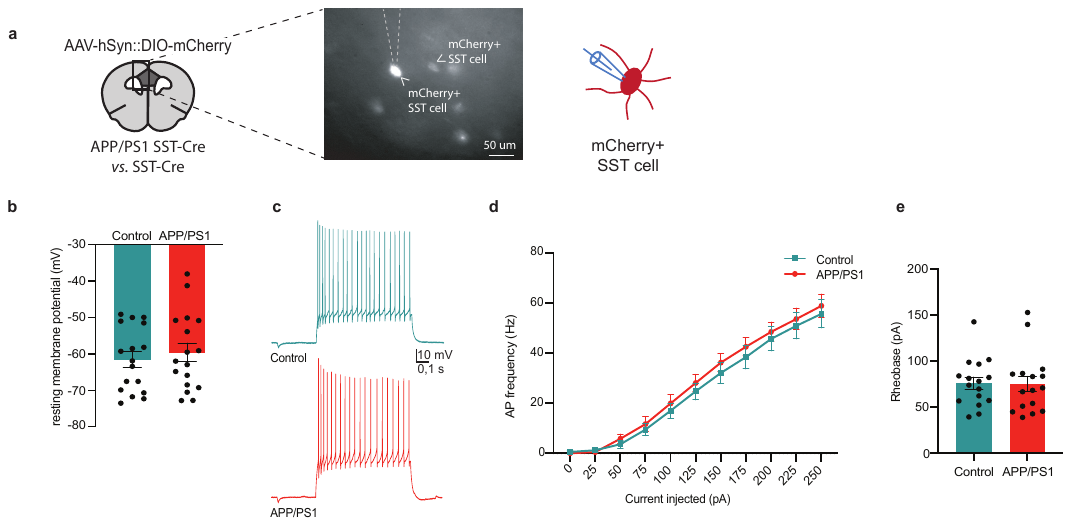

**Sup. Fig. 2:** SST cell excitability is unaltered in the mPFC of 20-week-old APP/PS1 mice. **a.** Schematic coronal brain section indicating the mPFC prelimbic region in dark grey, where AAV-hSyn::DIO-mCherry was microinjected and mCherry^+^ SST cells were recorded in APP/PS1 SST-Cre (APP/PS1) and SST-Cre (control) mice. Representative fluorescent image is depicted **b.** Resting membrane potential was unaltered in SST cells. Unpaired t-test: *t*_31_ = 0.73, *p* = 0.47, *n* = 17/16 cells, N = 5/7 control vs. APP/PS1 mice, respectively. **c.** Action potential (AP) firing of SST cells upon a depolarizing current step (250 pA) **d.** AP frequency in SST cells in response to 0-250 pA depolarizing current steps did not differ between genotypes. *Genotype x current* two-way repeated measures ANOVA *F*_(10,310)_ = 0.23, *p* = 0.99, *n* = 17/16 cells, N = 5/7 control vs. APP/PS1 mice. **e.** Rheobase was unchanged in SST cells Mann-Whitney test: *U* = 119, *p* = 0.75, *n* = 17/16 cells, N = 5/7 control vs. APP/PS1 mice, respectively. Graphs show mean ± s.e.m.

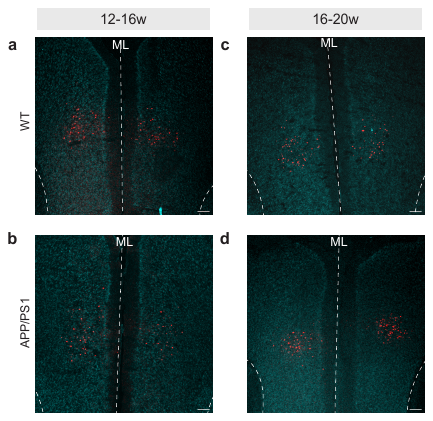

**Sup. Fig. 3:** Representative example of mCherry expression in the mPFC after CFC and 4TM treatment in APP/PS1 mice and WT controls, at 12-16w and 16-20w. A mixture of AAV-Fos::CreERT^2^ and AAV-Syn::DIO-mCherry was injected into the mPFC. Scale bar = 200 µm. ML= midline

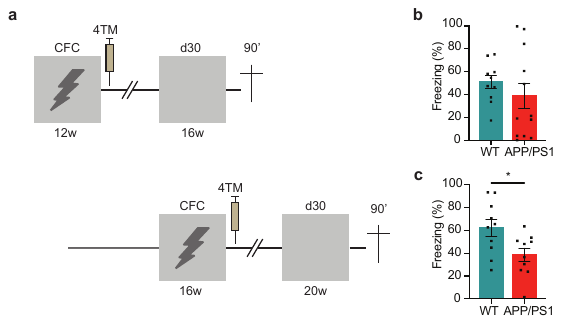

**Sup. Fig. 4**. APP/PS1 mice show remote memory impairment at 20, but not 16, weeks of age **a.** Experimental design. WT and APP/PS1 mice underwent CFC at 12 and 16 weeks of age, and memory retrieval 30 days later. Mice received an injection of 4-hydroxy tamoxifen (4TM) 2 hours after CFC to tag mPFC engram cells. **b.** At 16 weeks old, APP/PS1 mice did not differ in freezing levels compared to WT controls. Mann Whitney test: *U* = 46, *p* = 0.38, WT (*n* = 10), APP/PS1 (*n* = 12). **c.** At 20 weeks old, APP/PS1 mice show reduced freezing levels compared to WT controls. Unpaired *t*-test: *t*_18_ = 2.50, **p* = 0.022, WT (*n* = 10), APP/PS1 (*n* = 10). Graphs show mean ± s.e.m.

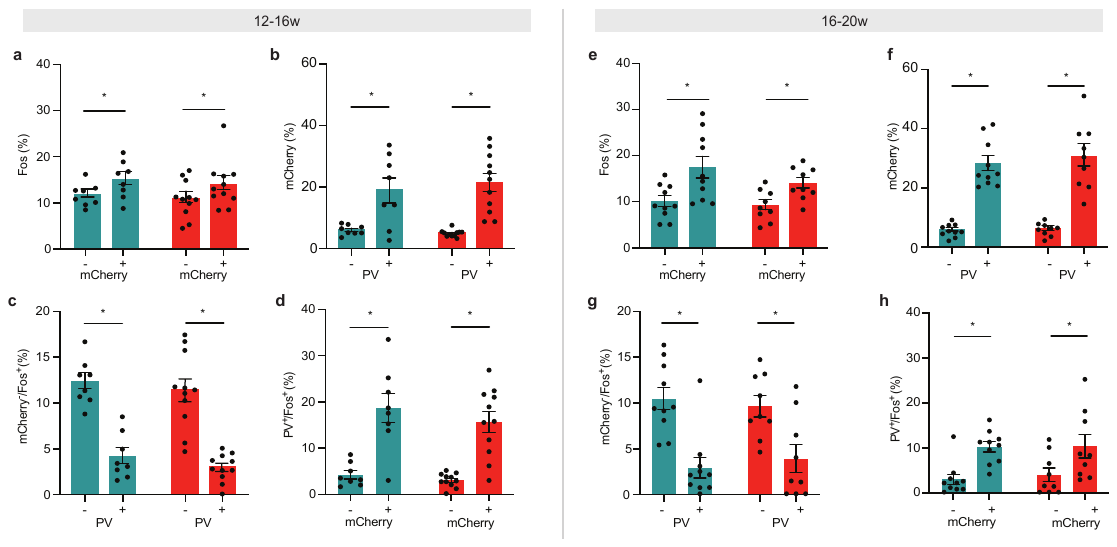

**sFig. 5: Size and reactivation of the mPFC engram ensemble, as well as PV interneuron (re)activation, are unaffected in APP/PS1 mice. a.** Fos colocalization with mCherry^+^ cells (mCherry^+^/Nissl^+^) was enhanced compared to mCherry^-^ cells (mCherry^-^/Nissl^+^) in WT and APP/PS1 mice. Two-way repeated measure ANOVA *cell population*: F_(1,17)_ = 16.3, **p* = 0.0009. Post-hoc Bonferroni test: Control **p* = 0.028; APP/PS1 **p* = 0.016. **b.** Percentage of PV^+^ cells in the mCherry^+^ population were higher than in the mCherry^-^ population for both genotypes. Two-way repeated measure ANOVA *cell population: F*_(1,17)_ = 31.70, **p* < 0.0001. Post-hoc Bonferroni test: WT **p* = 0.010; APP/PS1 **p* = 0.0003. **c.** Percentage of Fos^+^ cells were higher in the mCherry^-^/PV^-^ than mCherry^-^/PV^+^ population. Two-way repeated measure ANOVA *cell population*: *F*_(1,17)_ = 179.30, **p* <0.0001; Post-hoc Bonferroni test: WT **p* <0.0001; APP/PS1 **p* <0.0001. **d**. Percentage of Fos^+^ cells were higher in the PV^+^/mCherry^+^ population compared to the PV^+^/mCherry^-^ population in APP/PS1 and WT mice. Two-way repeated measure ANOVA *cell population: F*_(1,17)_ = 50.57, **p* < 0.0001; Post-hoc Bonferroni test: WT **p* = 0.0002; APP/PS1 **p* = 0.0002. **e**. Fos colocalization with mCherry^+^ cells (mCherry^+^/Nissl^+^) was enhanced compared to mCherry^-^ cells (mCherry^-^/Nissl^+^) in WT and APP/PS1 mice. Two-way repeated measure ANOVA *cell population*: *F*_(1,17)_ = 56.41, **p* < 0.0001. Post-hoc Bonferroni test: WT **p* < 0.0001; APP/PS1 **p* = 0.002. **f**. Percentage of PV^+^ cells in the mCherry^+^ population were higher than in the mCherry^-^ population for both genotypes. Two-way repeated measure ANOVA *cell population: F*_(1,17)_ = 82.15, **p* < 0.0001. Post-hoc Bonferroni test: WT control **p* < 0.0001; APP/PS1 **p* < 0.0001. **g**. Percentage of Fos^+^ cells were higher in the mCherry^-^/PV^-^ than mCherry^-^/PV^+^ population. Two-way repeated measure ANOVA *cell population*: *F*_(1,17)_ = 39.15, p <0.0001; Post-hoc Bonferroni test: WT **p* = 0.0001; APP/PS1 **p* = 0.004. **h**. Percentage of Fos^+^ cells were higher in the PV^+^/mCherry^+^ population compared to the PV^+^/mCherry^-^ population in APP/PS1 and WT mice. Two-way repeated measure ANOVA *cell population: F*_(1,17)_ = 17.38, **p* = 0.001. Post-hoc Bonferroni test: WT **p* = 0.010; APP/PS1 **p* = 0.031. Graphs show mean ± s.e.m.

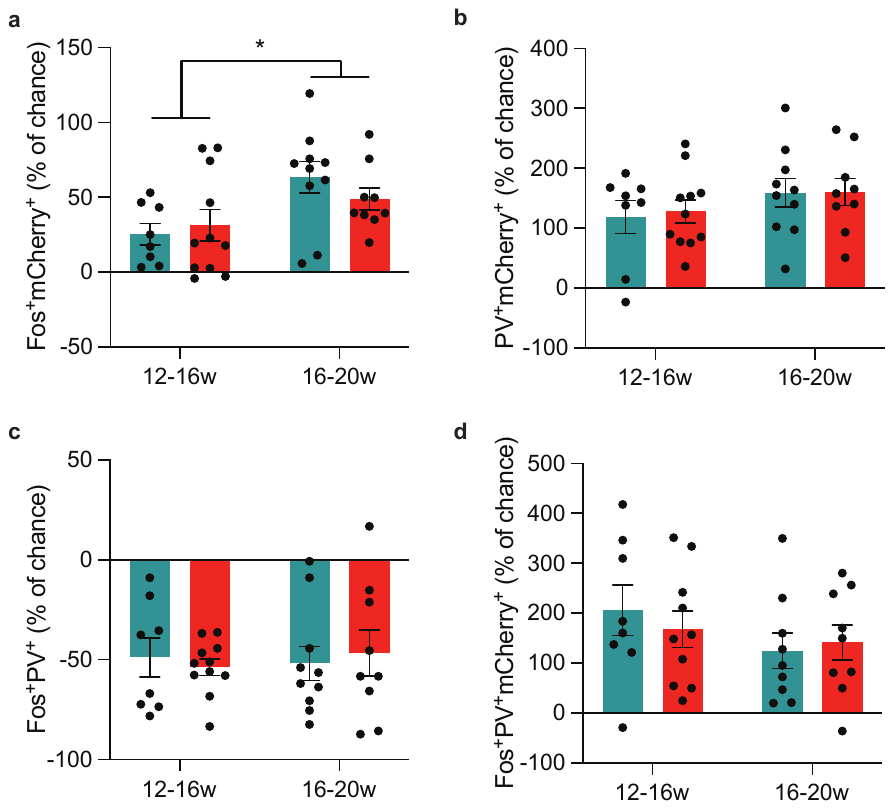

**sFig. 6: Increased reactivation of engram cells in 20- vs. 16-week-old WT and APP/PS1 mice. a**. Both genotypes showed increased overlap of Fos and mCherry at 16-20w compared to 12-16w. Two-way ANOVA *age groups F*_(1,34)_ = 8.57, *p* = 0.006. **b.** Overlap between PV and mCherry did not differ between genotypes or age groups. Two-way ANOVA *age groups F*_(1,34)_ = 2.45, *p* = 0.13. **c.** Overlap between Fos and PV did not differ between genotypes or age groups. Two-way ANOVA *age groups F*_(1,34)_ = 0.05, *p* = 0.82. **d.** Fos, PV, and mCherry did not differ between genotypes or age groups. Two-way ANOVA *age groups F*_(1,32)_ = 1.85, *p* = 0.18.

Supplemental tables

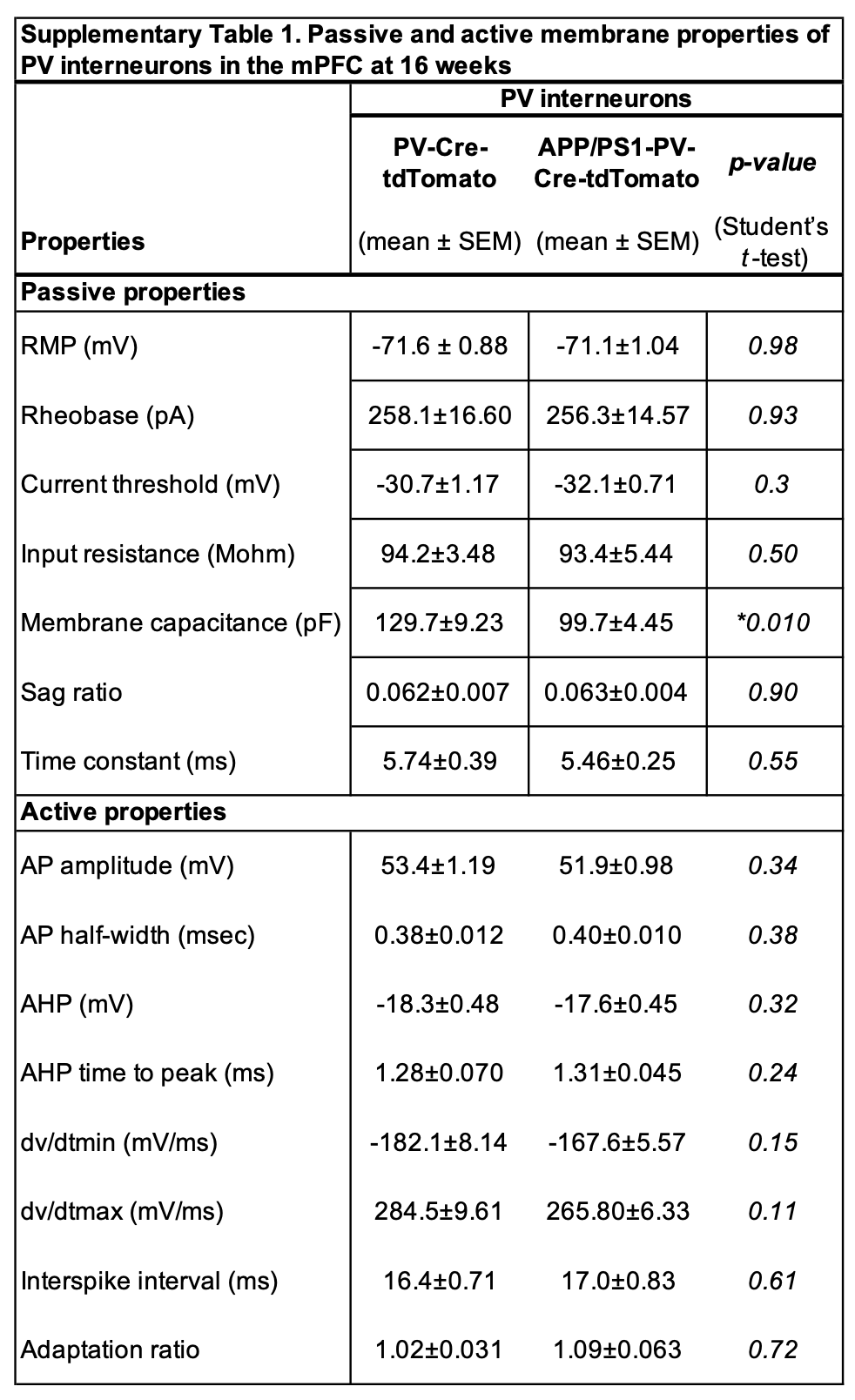

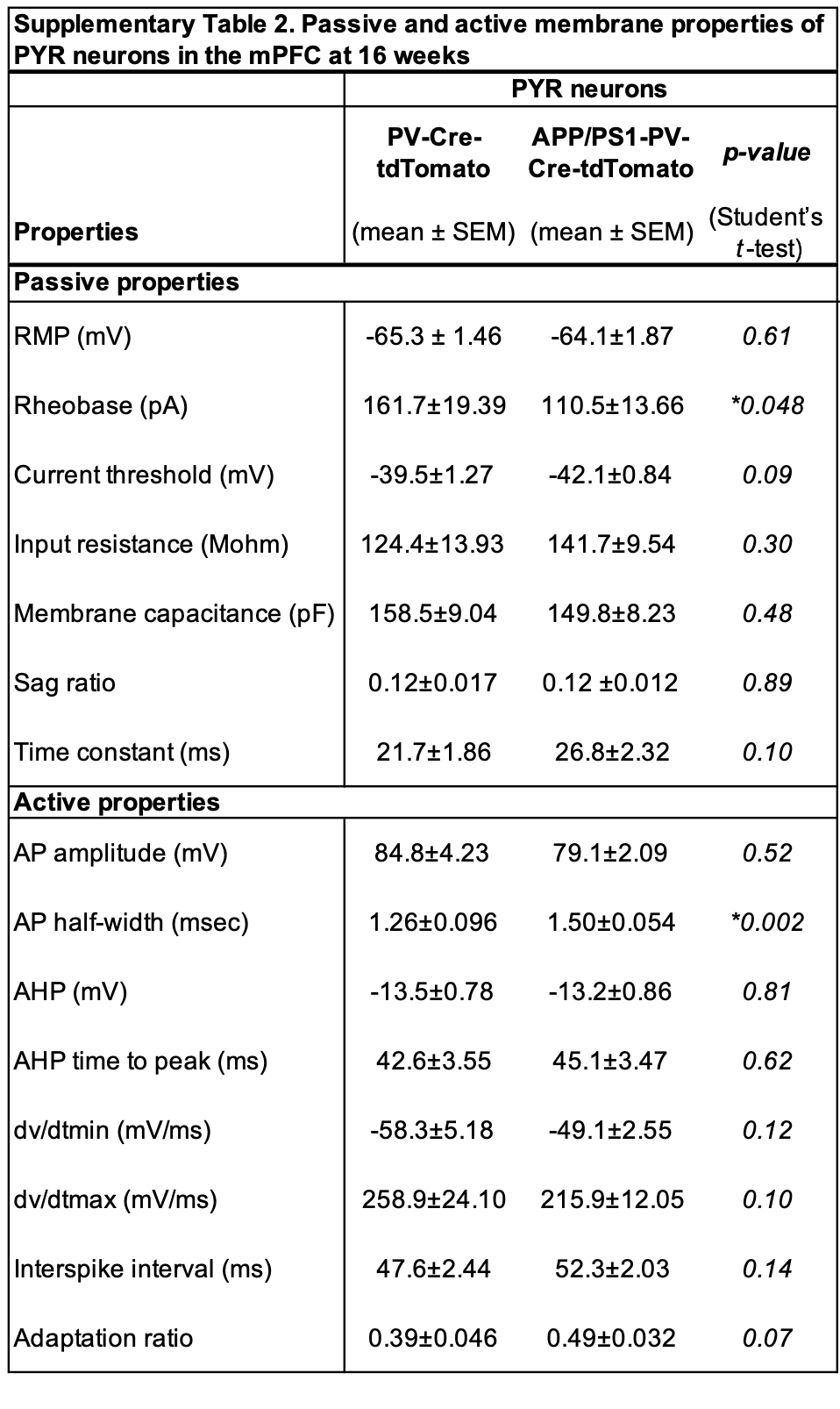

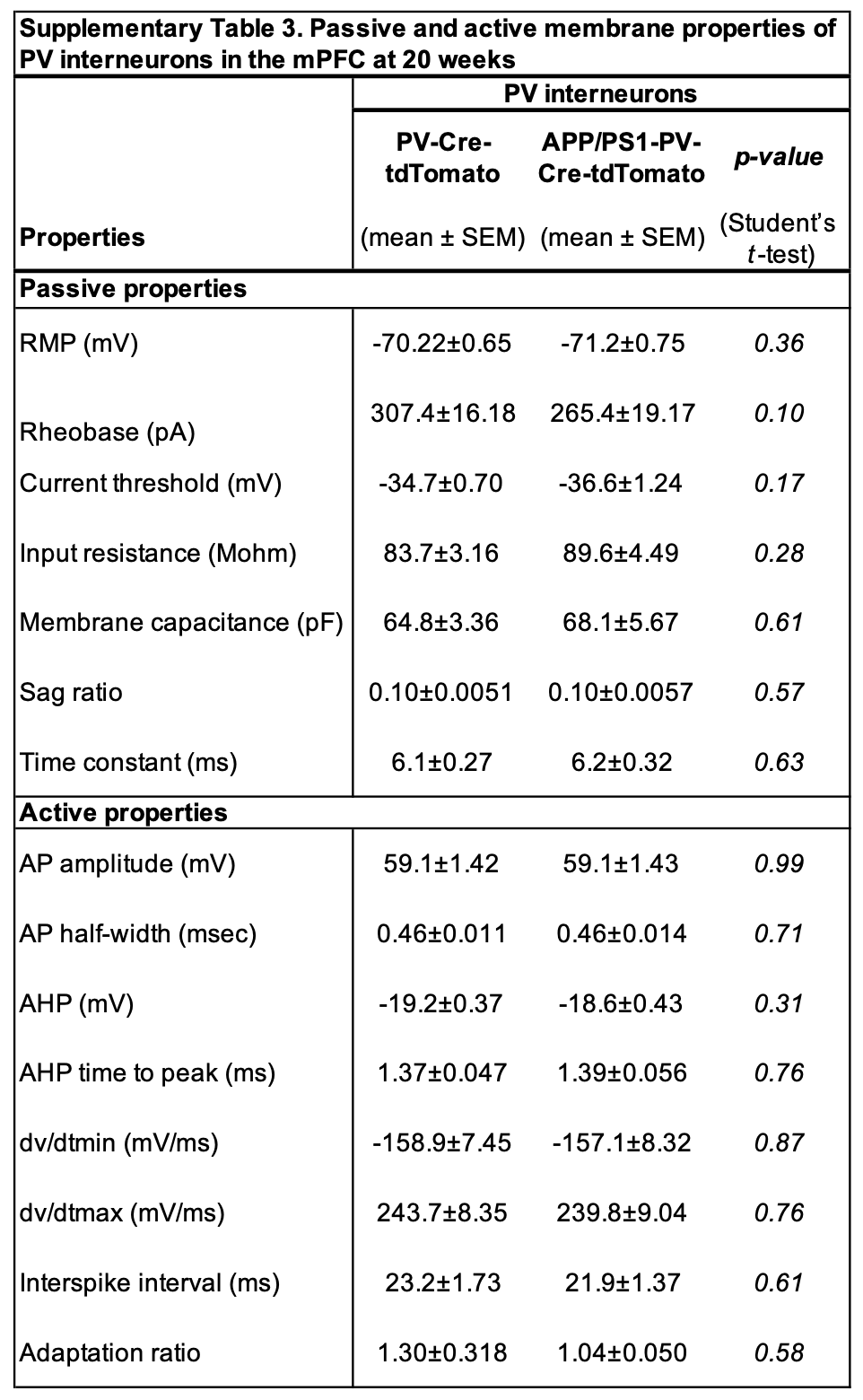

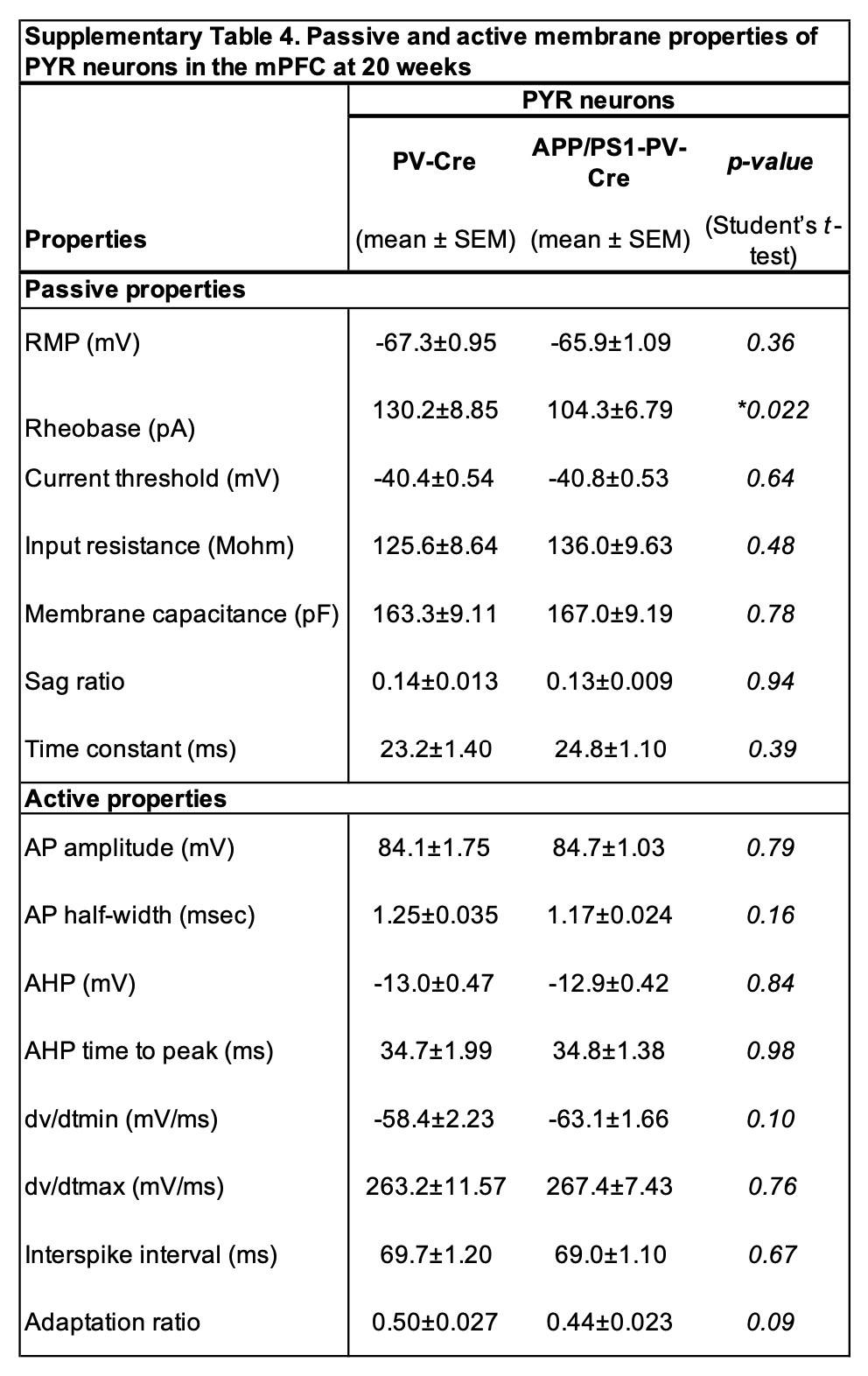

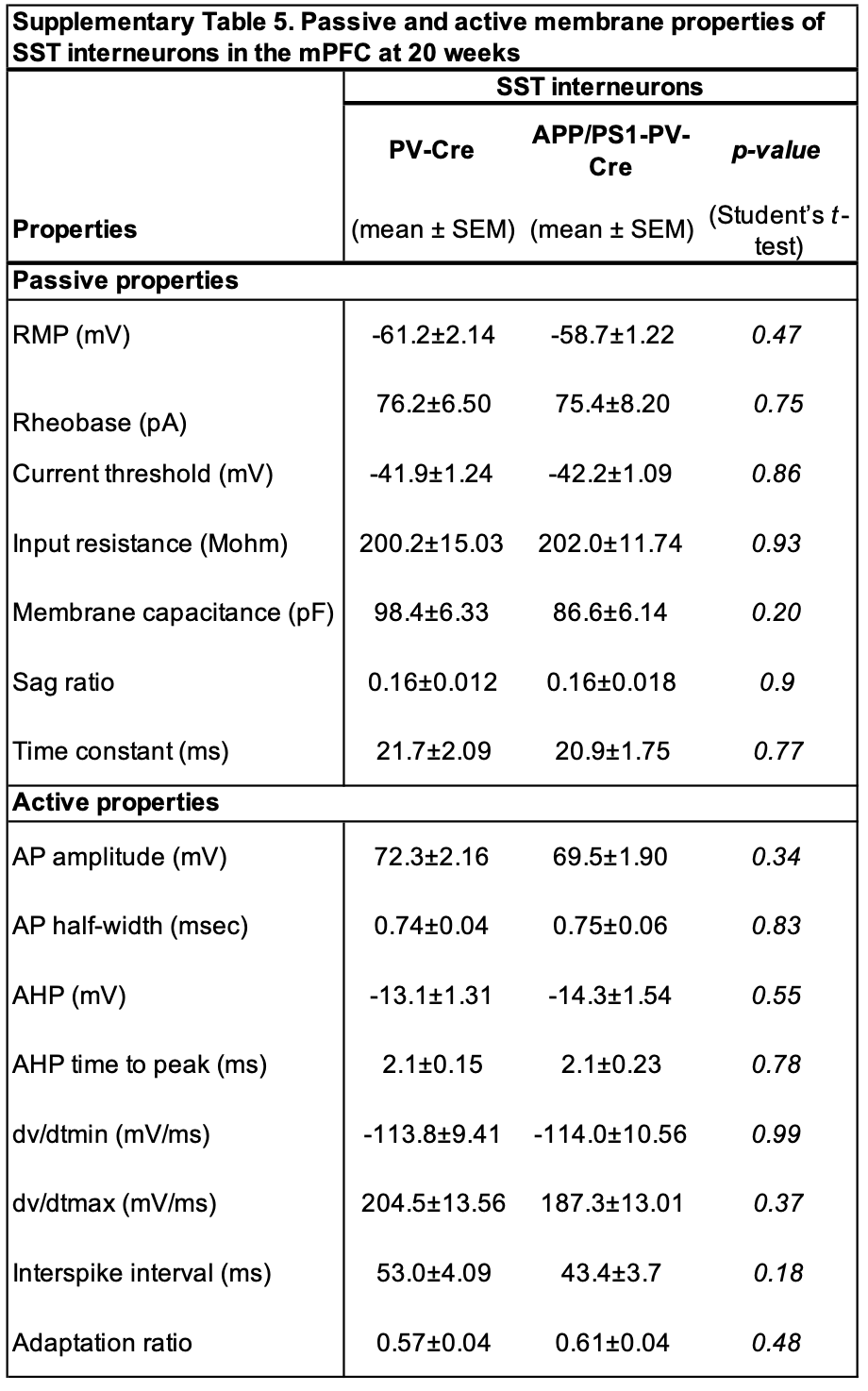
